# No detectable signal for ongoing genetic recombination in SARS-CoV-2

**DOI:** 10.1101/2020.12.15.422866

**Authors:** Damien Richard, Christopher J. Owen, Lucy van Dorp, François Balloux

**Affiliations:** UCL Genetics Institute, University College London, UK; Institute of Child Health, University College London, UK

## Abstract

The COVID-19 pandemic has led to an unprecedented global sequencing effort of its viral agent SARS-CoV-2. The first whole genome assembly of SARS-CoV-2 was published on January 5 2020. Since then, over 150,000 high-quality SARS-CoV-2 genomes have been made available. This large genomic resource has allowed tracing of the emergence and spread of mutations and phylogenetic reconstruction of SARS-CoV-2 lineages in near real time. Though, whether SARS-CoV-2 undergoes genetic recombination has been largely overlooked to date. Recombination-mediated rearrangement of variants that arose independently can be of major evolutionary importance. Moreover, the absence of recombination is a key assumption behind the application of phylogenetic inference methods. Here, we analyse the extant genomic diversity of SARS-CoV-2 and show that, to date, there is no detectable hallmark of recombination. We assess our detection power using simulations and validate our method on the related MERS-CoV for which we report evidence for widespread genetic recombination.

## Introduction

Genetic recombination is widely recognised as an important force in evolution, as it allows for the combination, within a single genome, of variants that arose independently in different genetic backgrounds [1]. Viruses are no exception to this pattern [2]. For example, recombination between its eight genomic segments (reassortment) is the fundamental mechanism behind the emergence of pandemic influenza A strains [3]. Recombination is also key for many viruses to generate new antigenic combinations that allow host immune systems evasion [4]. Moreover, the absence of recombination is also a prerequisite for phylogenetic inference [5]. Indeed, phylogenetic trees are limited to represent a single realisation of the past demography of the samples analysed. In the presence of genetic recombination, different regions of the sequence under scrutiny will support different evolutionary histories for the samples analysed, and hence result in conflicts in the topology of the phylogenetic reconstruction [5]. Not accounting for recombination can also lead to false positive detection of sites undergoing positive selection [6, 7].

Repositories of Severe acute respiratory syndrome coronavirus 2 (SARS-CoV-2) genomes have grown at an unprecedented pace, with over 150,000 high-quality complete genome assemblies currently available on the Global Initiative on Sharing All Influenza Data (GISAID) repository as of 17/11/2020 [8, 9]. This allows for the near-real-time monitoring of the emergence and spread of novel mutations [10, 11] and the description of emerging lineages [12, 13]. Most of these studies rely on phylogenetic reconstructions of genome-wide Single Nucleotide Polymorphisms (SNPs), and implicitly assume the absence of pervasive genetic recombination in SARS-CoV-2. So far there has been limited effort to assess the extent of ongoing recombination in SARS-CoV-2 ([14, 15], https://observablehq.com/@spond/linkage-disequilibirum-in-sars-cov-2) despite its potential relevance to understanding the duration and propensity of co-infections in host.

Conflicts (incongruence) between phylogenies inferred from different genome segments can be indicative of recombination. Some of the numerous methods developed to detect genetic recombination rely on this concept [16–19]. The so-called “compatibility test” checks if all four combinations of alleles of a pair of biallelic sites (00, 01, 10, 11) are present among the sequences. More refined methods relying on this principle have been developed including the pairwise homoplasy index (PHI) [20]. Recombination also has the effect of decorrelating allele frequencies, with this effect increasing with physical distance along the genome. In a population undergoing frequent recombination, this causes linkage disequilibrium of alleles to decay with physical distance on the sequence [4]. The *r*^2^ metric [21] is commonly used to measure linkage disequilibrium [22].

In this work, we aimed to detect signals of genetic recombination within the SARS-CoV-2 global population. We assembled a curated alignment of 6,546 available SARS-CoV-2 genomes enriched for those collected more recently to maximise genetic diversity in the dataset, and hence our ability to detect recombination. We applied two different statistical methods for the detection of genetic recombination and assessed their power using bespoke simulations. We validate our methodology on the related *Betacoronavirus* Middle East respiratory syndrome-related coronavirus (MERS-CoV) [23] responsible for the MERS outbreaks beginning in 2012, for which we find evidence for recombination, consistent with previous reports [24]. Our results do not identify detectable evidence for recombination in the SARS-CoV-2 population as of September 2020.

## Results

### No signal of recombination in SARS-CoV-2

We compiled an alignment of 6,546 SARS-CoV-2 isolates sampled across six continental regions. In order to maximize our detection power, whilst keeping the dataset computationally manageable, we chose to restrict our analysis to the most recently collected genomes during the month of September 2020. This is expected to maximise the genetic diversity in the dataset, and hence increase power to detect recombination events. We detect over 4,000 polymorphic positions following masking of sites putatively suggested as artefactual ([25] https://github.com/W-L/ProblematicSites_SARS-CoV2/blob/master/problematic_sites_sarsCov2.vcf, accessed 29/10/2020). These include a large fraction of homoplasies (29.6%), thought to largely be induced by host immune system RNA editing [15, 26, 27].

A PHI test applied to the SARS-CoV-2 alignment reported a *p*-value of 0.78 suggesting no signal of recombination. Consistently, LD decay regression coefficients and R squared statistics did not fall outside of the distributions obtained after randomly permuting genome coordinates (Figure 1).

**Figure 1:**
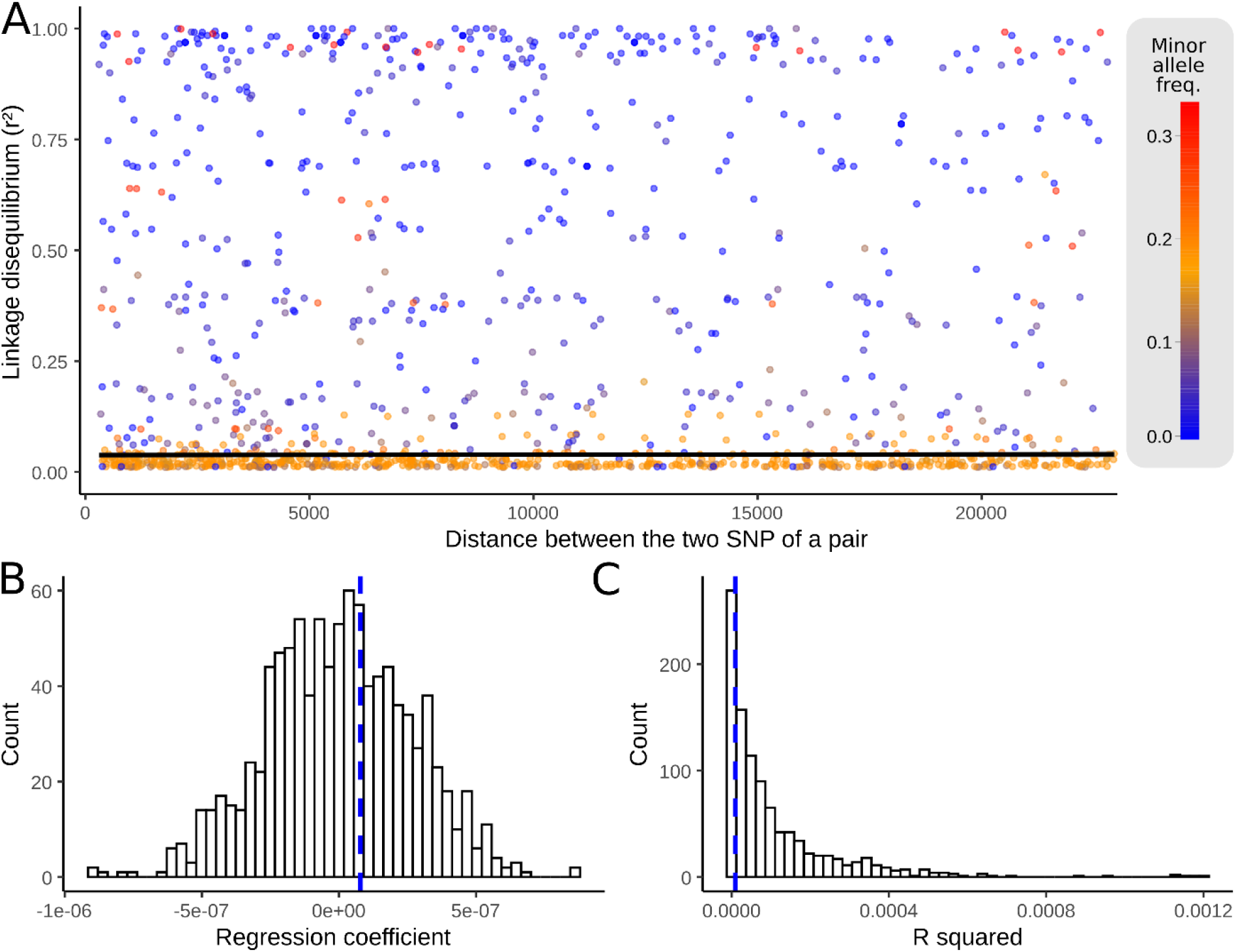
Linkage disequilibrium (*r*^2^) as a function of physical distance on the SARS-CoV-2 genome. (A) Linkage disequilibrium measured by *r^2^* (y-axis) for all pairs of SNPs represented as a function of the genetic distance separating the SNPs of each pair. Black line: fitted linear model (regression coefficient: 7.84e-8; R-squared: 9.33e-6). (B) Distribution of regression coefficients of the linear models obtained following consideration of 1000 position permuted datasets. Blue dashed line: value obtained for the true SARS-CoV-2 alignment. (C) Distribution of the R squared values of the linear models of 1000 position permuted datasets. Blue dashed line: value of the SARS-CoV-2 true alignment.

To test our ability to detect low levels of recombination, we simulated alignments with levels of genetic diversity matching that observed in the true SARS-CoV-2 alignment, but using varying recombination rates. We detected recombination in 100% of the datasets simulated with 3e-3 recombination events per genome per viral replication (60% for a rate of 3e-4, Supplementary Table S1). This low detection power is linked to the high homogeneity of the SARS-CoV-2 population, reflected by the mean pairwise distance of 19.4 (95%HPD 3-30) SNPs in the alignment analysed.

In addition, we searched the global SARS-CoV-2 phylogeny for isolates displaying root-to-tip distances in the upper 5% quartile of the distribution. These may offer some of the best candidates for isolates having experienced recombination, which is expected to increase terminal branch length. We detected 24 phylogenetic outliers which grouped into 13 phylogenetic clades. Localisation of their mutations and those of their phylogenetic neighbours in matrices did not support a recombinant origin. Indeed, rather than displaying syntenic groups of private mutations that would suggest a recombination-mediated origin, the 24 outliers mostly displayed a randomly distributed excess of mutations (Supplementary Figures S1-S13). Our results therefore suggest that recombination in SARS-CoV-2 is either absent, or occurring at a rate too low relative to mutation to be detectable under the genetic diversity charactering the SARS-CoV-2 population at this stage.

### Recombination occurs in MERS-CoV

To validate our approach, we applied the same method to an alignment of MERS-CoV, a related *Betacoronavirus* thought to be widely recombining [24, 28]. Counter to observations for SARS-CoV-2, the MERS-CoV dataset yielded detectable evidence of recombination. Beside the decay of the r^2^ ~ distance regression slope, values of both R-squared and the regression coefficient largely fall outside of the distributions of the same parameters of the permuted datasets (Figure 2). The PHI test reported a p-value <1e-12. Of note, these tests were repeated after discarding C to T mutations, mostly caused by the host immune RNA editing systems, that might produce an artefactual signal of recombination. Tests on the pruned alignment still provide evidence of significant signals of recombination in MERS-CoV (PHI test p-value <1e-12 and significant decay of r^2^, Supplementary Figure S14).

**Figure 2:**
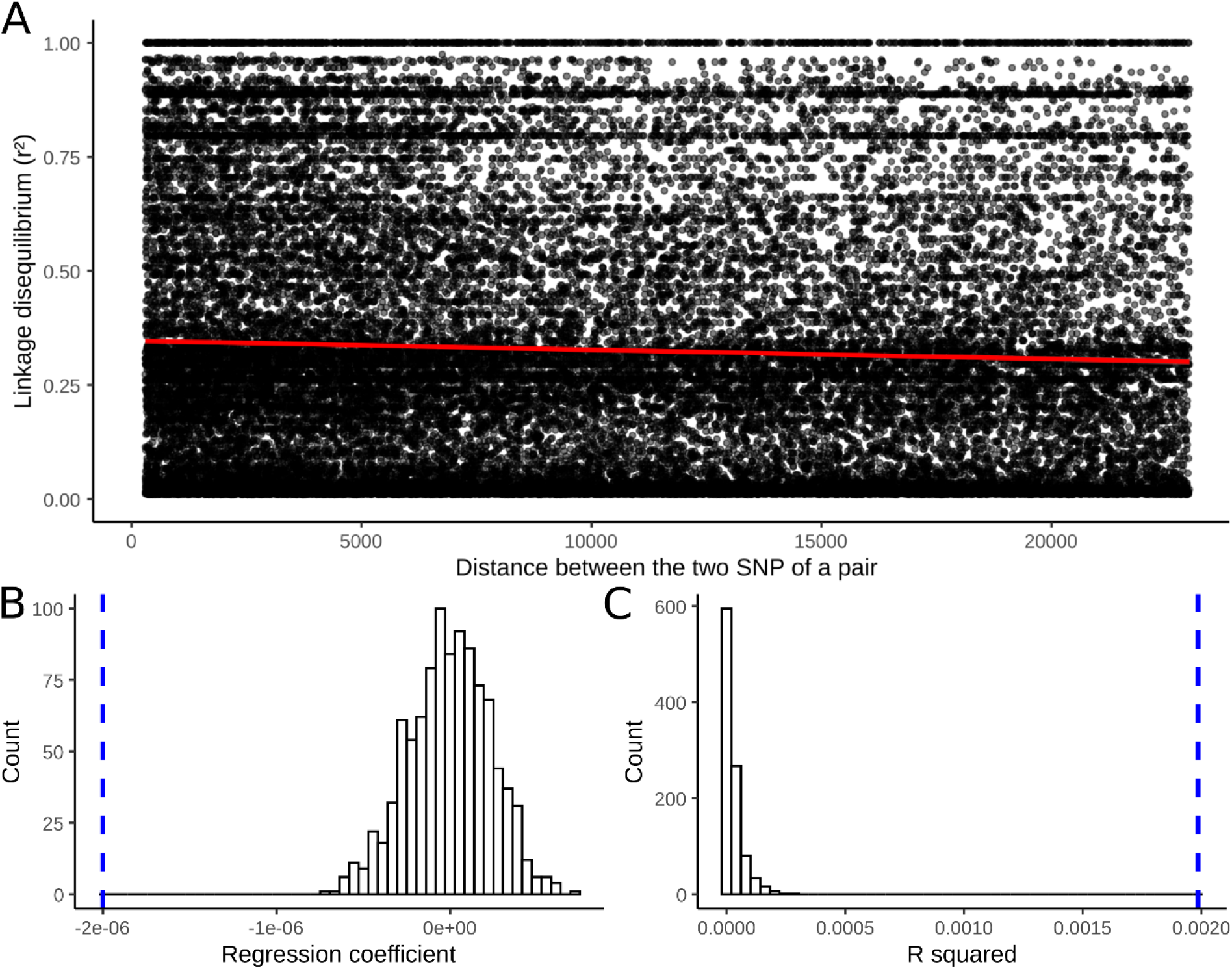
Linkage disequilibrium (*r*^2^) as a function of physical distance on the MERS-CoV genomes. Pairs comprising SNP differing by a frequency >=0.1 have been discarded, lowering the number of pairs from 261,090 to 212,342. (A) Linkage disequilibrium (y-axis) for all pairs of SNPs is represented as a function of the distance separating the SNPs of each pair. Red line: fitted linear model (regression coefficient: −2.00e-6; R-squared: 1.99e-3). (B) Distribution of the regression coefficients of the linear models obtained following consideration of 1000 position permuted datasets. Blue dashed line: value of the MERS-CoV true alignment. (C) Distribution of the R squared values of the linear models of 1000 position permuted datasets. Blue dashed line: value of the MERS-CoV true alignment.

## Discussion

In this study, we analyzed a dataset of 6,546 SARS-CoV-2 assemblies sampled during the month of September 2020 across six continental regions. Applying two distinct detection methods, we did not find signal of recombination among the SARS-CoV-2 population tested. Conversely, we detected evidence of recombination in SARS-CoV-2-like simulated recombining datasets as well as in a set of 459 MERS-CoV coronaviruses assemblies, known for being prone to recombination.

*A priori* it is highly plausible that SARS-CoV-2 has the potential to recombine [28]. Recombination has been suggested to be common in coronaviruses, including both for human [29–33] and animal [34–36] associated lineages, as inferred from genomic approaches [37], observed in cell culture [38, 39] and *in vivo* [40]. It has been claimed that all of the human epidemic coronaviruses: SARS-CoV-1 [41–43], MERS-CoV [44], and SARS-CoV-2 [45–47] may have evolved through recombination events leading to some genome mosaicism, particular over receptor binding regions.

At this stage we do not detect a genetic hallmark for recombination in SARS-CoV-2. This does not necessarily imply that SARS-CoV-2 lacks the ability to recombine. For genetic recombination to leave a measurable signal in the genetic data, there needs to be sufficient genetic differentiation between the recombining viruses. Given the low intra-host genetic diversity of epidemic viruses such as SARS-CoV-2, this requires mixed infections (i.e. coinfection of the same host by distinct SARS-CoV-2 lineages). While such events are expected to be rare, there have been reports of mixed infections [48]. Though, given the limited genetic diversity of SARS-CoV-2 strains currently in circulation, even mixed infections may often not involve sufficiently differentiated strains to leave a detectable signal following a recombination event. The recent host jump into humans of SARS-CoV-2, most likely through a single transmission to humans from an unknown animal reservoir, created an essentially genetically invariant viral population, with genetic diversity building up through the accumulation of mutations since the begininning of the pandemic. The genomic diversity of SARS-CoV-2 is still far below its mutation-drift equilibrium and remains very low at this stage [10]. As a result, putative recombination events would only be supported by a limited number of SNPs, and would require high detection sensitivity to be identified.

In contrast, we detected recombination in MERS-CoV despite the far smaller sample size of the alignment analysed. Besides the higher genetic diversity of MERS-CoV at this stage, this may also be due to major epidemiological differences between MERS and COVID-19. MERS is mainly a disease of dromedary camels, with spillover events into humans [28]. In camels, the high prevalence of the disease and the mostly mild symptoms it causes is suggested to favour co-infection [24]. On the contrary, the severity of the symptoms in human lowers the probability of co-infection. The camel host, which is known to harbour other coronaviruses, could provide a hub of genetic diversity creation in MERS-CoV through recombination [49]. Human MERS-CoV infection was first documented in 2012, but it is thought the virus had been previously circulating for at least a few years in camels [50]. Our MERS-CoV alignment comprises samples spanning from 2012 to 2019. Mutations accumulating over this time-scale provide more diverse genetic markers that facilitate the detection of putative recombination events.

While we did not detect evidence of genetic recombination in SARS-CoV-2 to date, it remains of importance to repeat such analyses as the genetic diversity of the SARS-CoV-2 population will increase, and to consider its possible impact when conducting phylogenetics studies in the future.

## Materials and Methods

### SARS-CoV-2 dataset

All 6,546 SARS-CoV-2 high quality genomes (containing less than 5% of “N” and being >29,000 bp long) sampled during the month of September 2020 available on GISAID (as of October 15^th^ 2020) were downloaded and profile aligned to the Wuhan-Hu-1 reference genome (GenBank accession MN908947; GISAID ID EPI_ISL_402125) using MAFFT v7.471 [51]. A full list of acknowledgements together with submitting and originating laboratories is provided in Supplementary Table S2. SNPs flagged as putative sequencing errors were discarded (https://github.com/W-L/ProblematicSites_SARS-CoV2/blob/master/problematic_sites_sarsCov2.vcf, accessed 29/10/2020). The final dataset comprised 4,199 SNPs and a mean pairwise SNP difference of 19.4 (95%HPD 3-30). Following construction of a maximum likelihood tree using IQTree Covid-release [52], 29.6% of SNPs were identified as homoplasic by HomoplasyFinder [53].

### MERS dataset

456 high-quality MERS-CoV genomes isolated from both camels and human were downloaded from the NCBI Virus database and profile aligned to the HCoV-EMC/2012 reference genome (GenBank accession NC_019843) using MAFFT v7.471 [51] (Supplementary Table S3). We detected 8,788 SNPs in the alignment, with a mean pairwise SNP count of 123.36 (95%HPD 18-234). Homoplasies were identified as described above and represented 12.4% of the polymorphic positions.

### Detection of recombination

Two recombination tests were performed on each dataset. First, a pairwise homoplasy index (PHI) test was used to detect recombination setting the number of permutations to 100 and the window size set to 300 bp with otherwise default parameters [20]. Additionally, we computed the linkage disequilibrium (*r*^2^) for all pairs of bi-allelic SNPs occurring in >=1% of the isolates using tomahawk (https://mklarqvist.github.io/tomahawk/). 90% of the SNP pairs grouped <= 23,000 nucleotides apart. Linkage disequilibrium estimation over distances larger than 23,000 rely on a few SNP pairs only, so we restricted the dataset to those 90% pairs. A linear model was fitted to the distribution of *r*^2^ values as a function of the distance separating the two SNPs in each pair. The regression coefficient of this linear model indicates whether linkage disequilibrium decays with physical distance or not. To formally test for the presence of recombination, we produced 1000 permuted datasets (randomly associating *r*^2^ values with distance values) and fitted a linear model to each one of the permuted datasets. We then assessed whether the real R-squared and regression coefficients values fell either inside or outside of the distributions of the parameters generated by the randomly permuted datasets. A limitation of the use of the *r*^2^ metrics as an estimator of linkage disequilibrium is its dependency on allele frequencies, causing a possible reduction in statistical power [54]. It has therefore been proposed to compute *r*^2^ only for pairs of SNPs that do not differ markedly in frequency in the studied population [55]. Discarding pairs of SNPs is suboptimal in the context of SARS-CoV-2’s already restricted genetic diversity. However, we still implemented this approach which led to similar results to tests on non-frequency filtered SNPs (Supplementary Figure S15).

We performed a third test for recombination in SARS-CoV-2 by focusing specifically on isolates that were flagged as phylogenetic outliers in the global phylogeny. Recombinant isolates are expected to be located at the tip of long terminal branches if there is phylogenetic incongruency between the mutations they carry. We applied TreeShrink to identify the accessions displaying root-to-tip distances in the upper 5% quartile (-q 0.05 parameter) of the root-to-tip distance distribution [56]. The mutations carried by those outliers were visualy compared to that of their neighbours in the phylogeny.

### Power of recombination detection

In order to characterise the statistical power of the recombination detection methods employed, we simulated in silico SARS-CoV-2 alignments using MSprime [57]. The simulated mutation rate was set to match that of the real dataset (Supplementary Figure S16). We generated datasets with numbers of recombination events per genome per viral replication of 0, 3e-7, 3e-6, 3e-5, 3e-4, 3e-3 and 3e-2 (ten replicates each). PHI tests and linkage disequilibrium decay tests were performed on those simulated datasets as described previously.

## Data and Code Availability

All analysed SARS-CoV-2 data is available on registration to GISAID with the accession IDS and acknowledgements provided in Table S2. MERS-CoV assemblies are freely available on NCBI with the included accessions provided in Table S3. Scripts used in this study are available at https://github.com/DamienFr/LD_SARS-CoV-2.

## Acknowledgements

D.R. is supported by a NIHR Precision AMR award. C.O is funded by a NERC-DTP studentship. L.v.D and F.B. acknowledge financial support from the Newton Fund UK-China NSFC initiative (grant MR/P007597/1) and the BBSRC (equipment grant BB/R01356X/1). L.v.D. is supported by a UCL Excellence Fellowship. We wish to particularly acknowledge all of the large number of contributing and submitting laboratories sharing SARS-CoV-2 assemblies via the GISAID platform, including the UK (COG-UK) consortium (a full list of consortium names and affiliations can be found at https://www.cogconsortium.uk). COG-UK is supported by funding from the Medical Research Council (MRC) part of UK Research & Innovation (UKRI), the National Institute of Health Research (NIHR) and Genome Research Limited, operating as the Wellcome Sanger Institute.

