## Supplementary File for "No detectable signal for ongoing genetic recombination in SARS-CoV-2"

Table S1: External Excel document. Number of simulations displaying (i) R squared and (ii) regression coefficient significantly different from the NULL permuted distribution (1,000 permutations), using alpha=0.01 and (iii) PHI tests reporting p-value < 0.01.

Table S2: External Excel document. Accessions of SARS-CoV-2 used in this study, including full acknowledgement of originating and submitting labs.

Table S3: External Excel document. NCBI accessions of MERS-CoV used in this study.

Figure S1

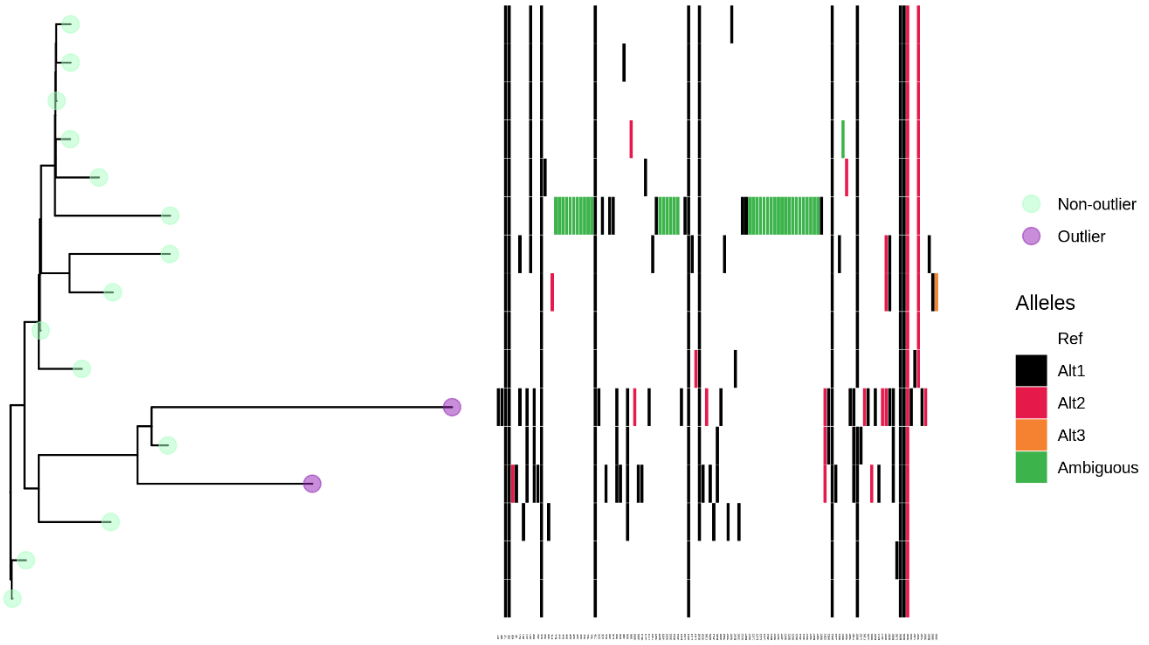


Figure S2

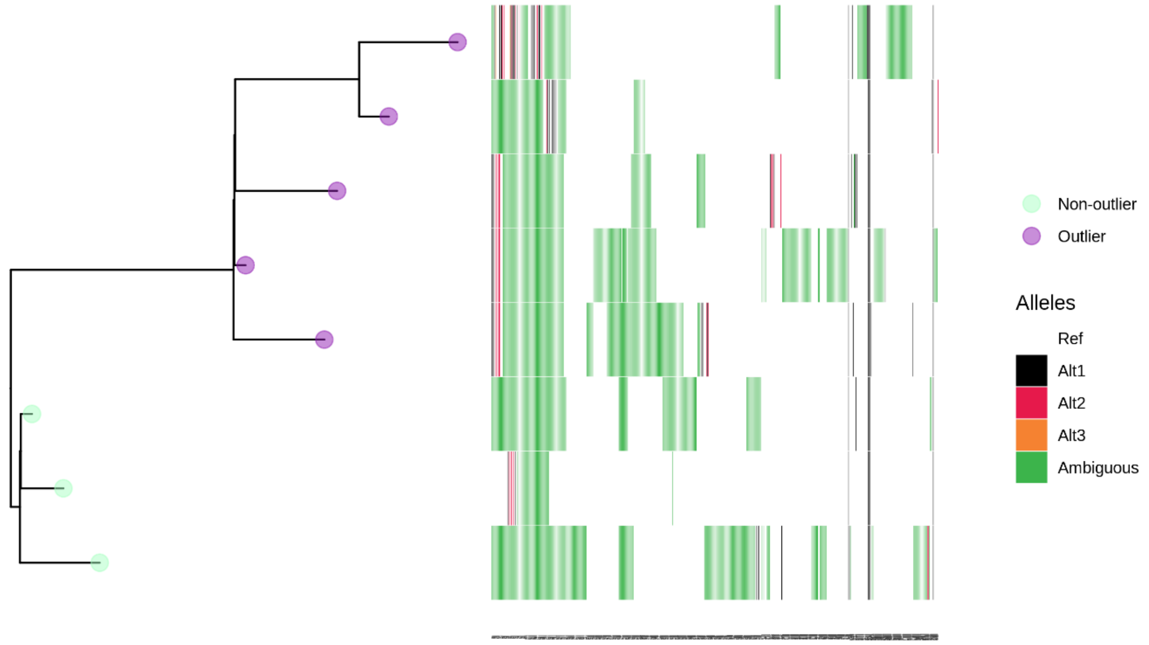

Figure S3

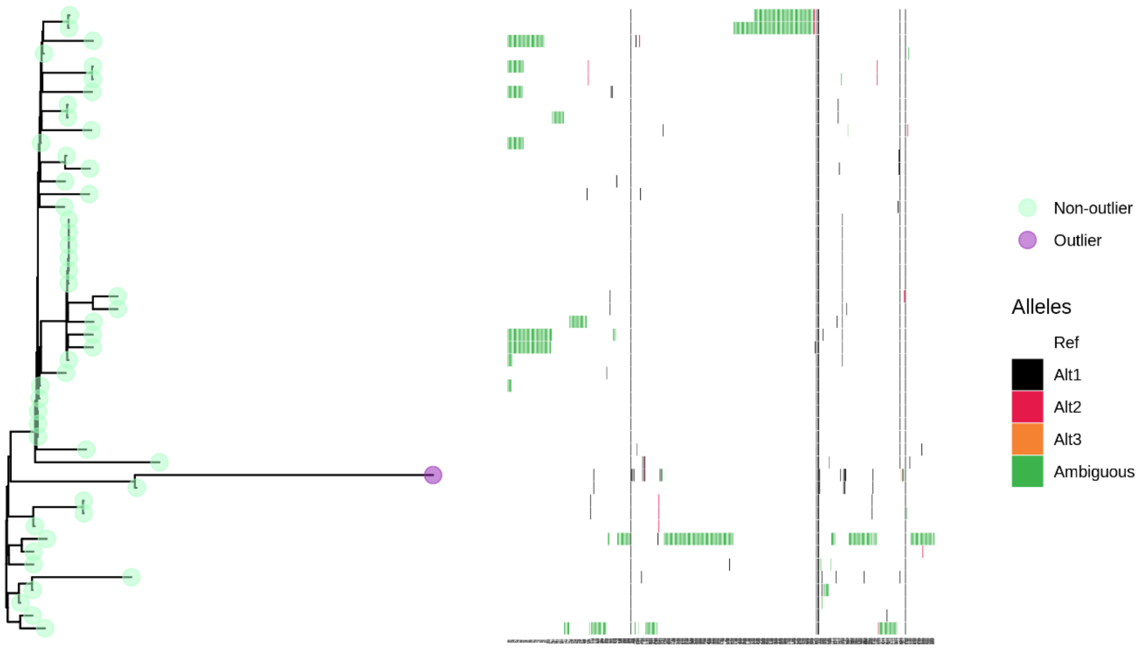


Figure S4

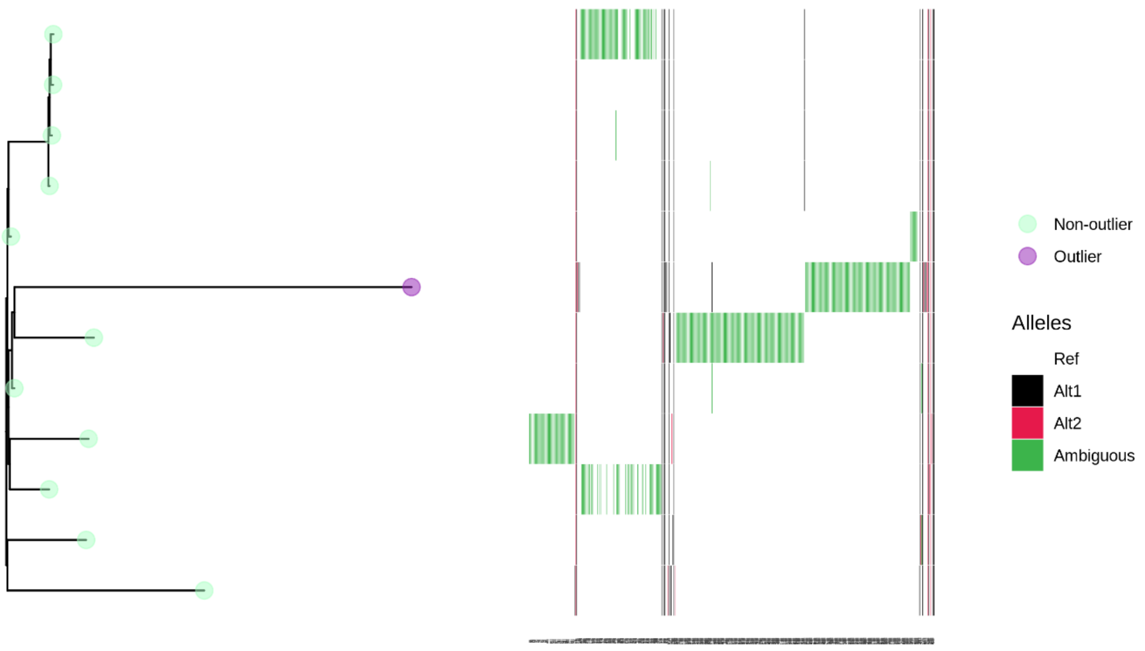

Figure S5

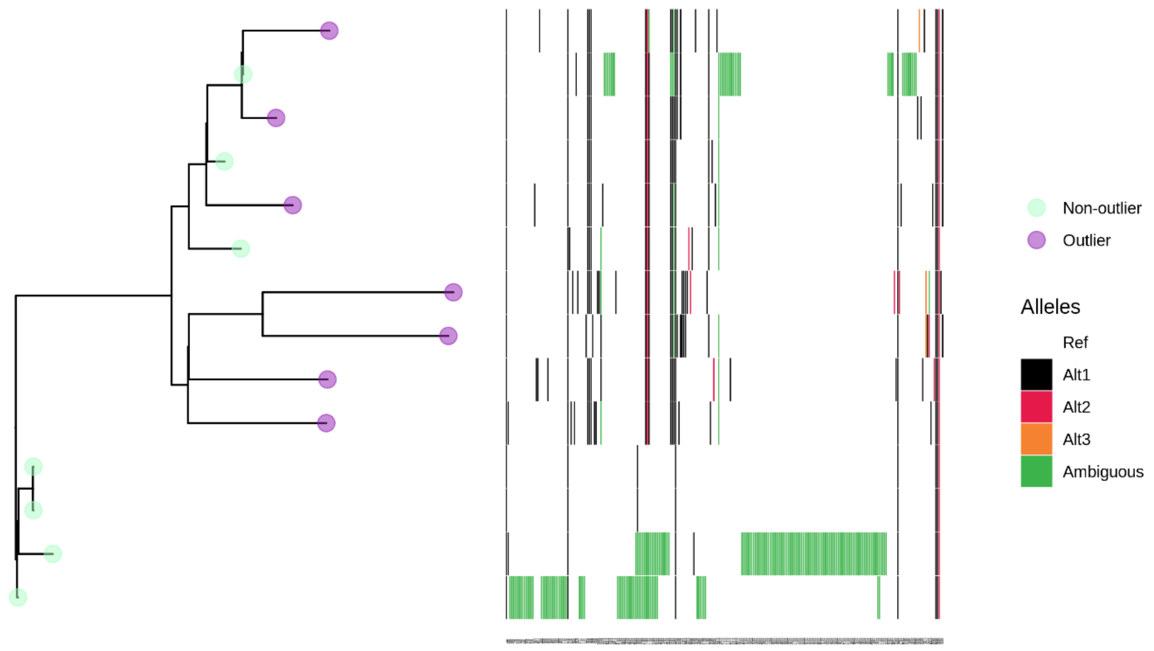


Figure S6

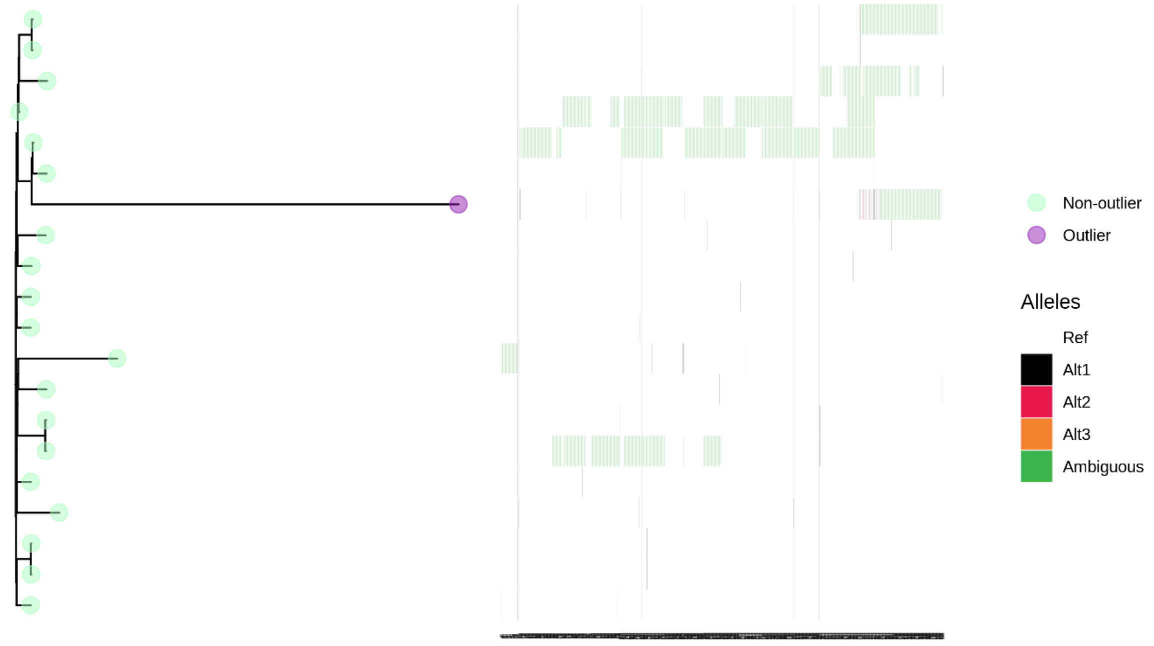

Figure S7

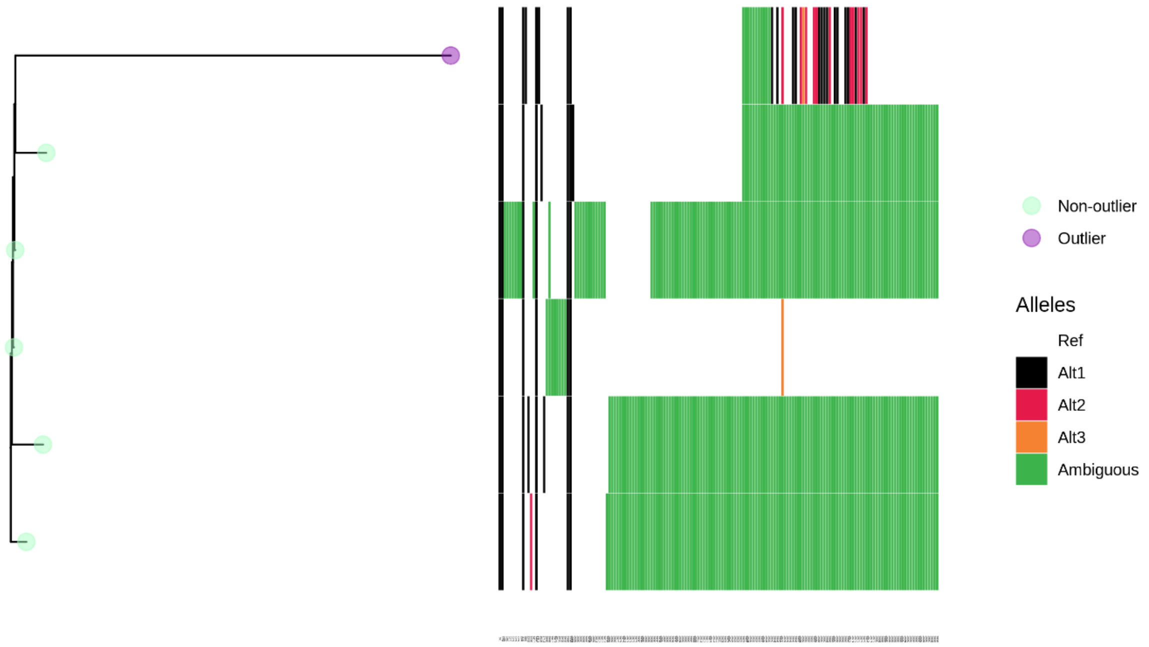


Figure S8

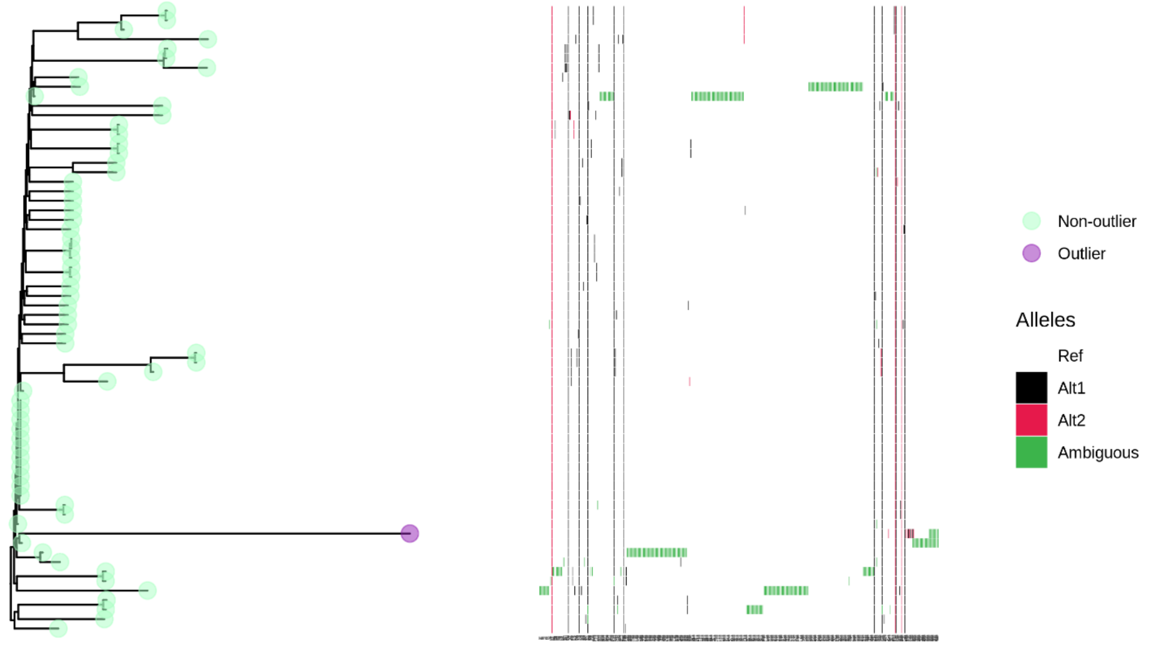

Figure S9

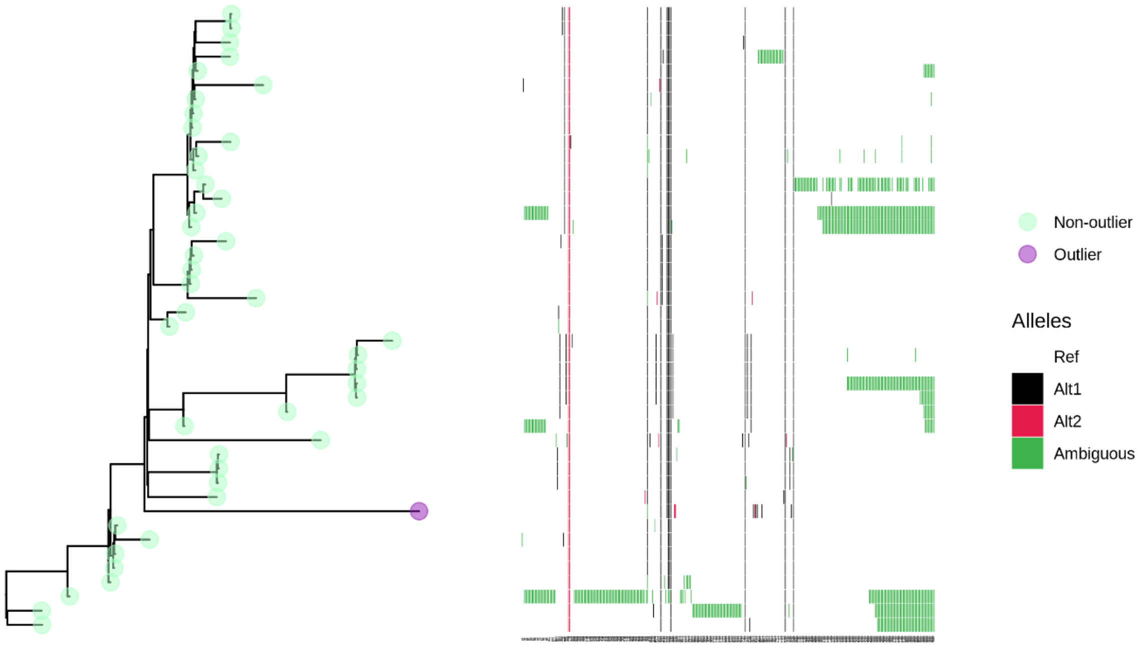


Figure S10

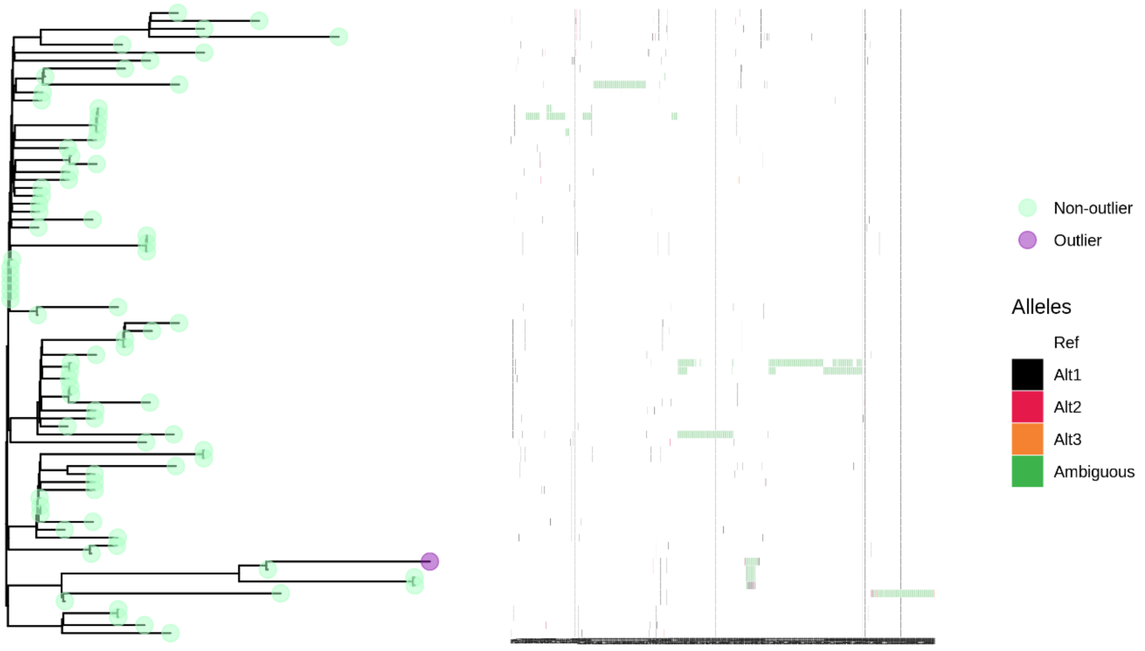

Figure S11

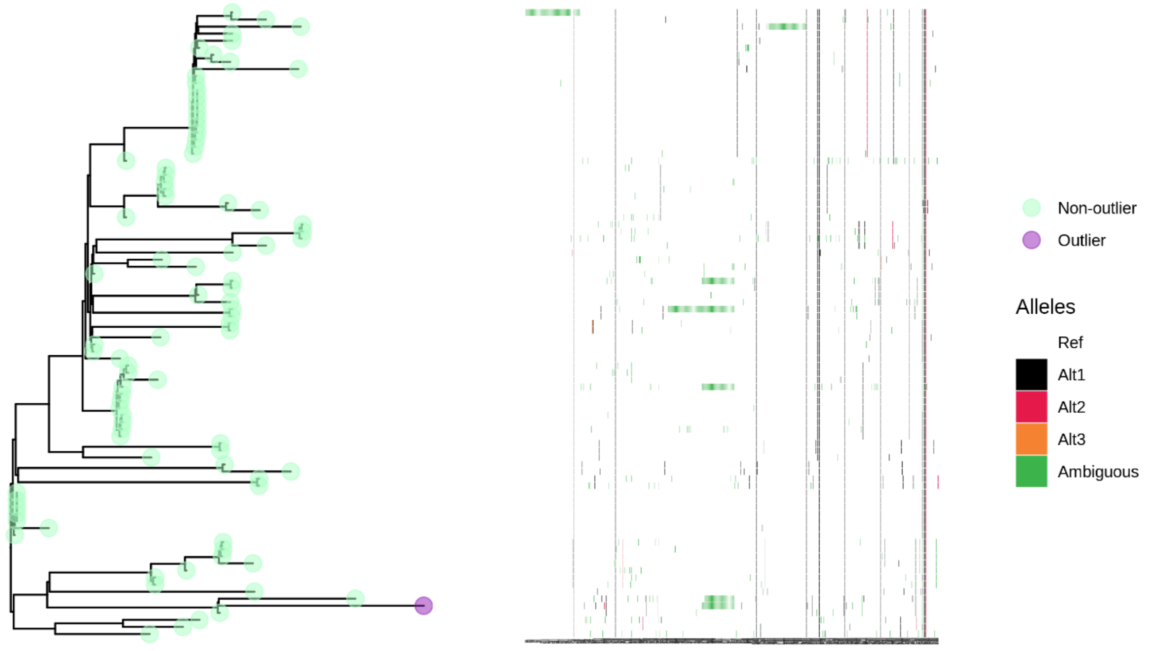


Figure S12

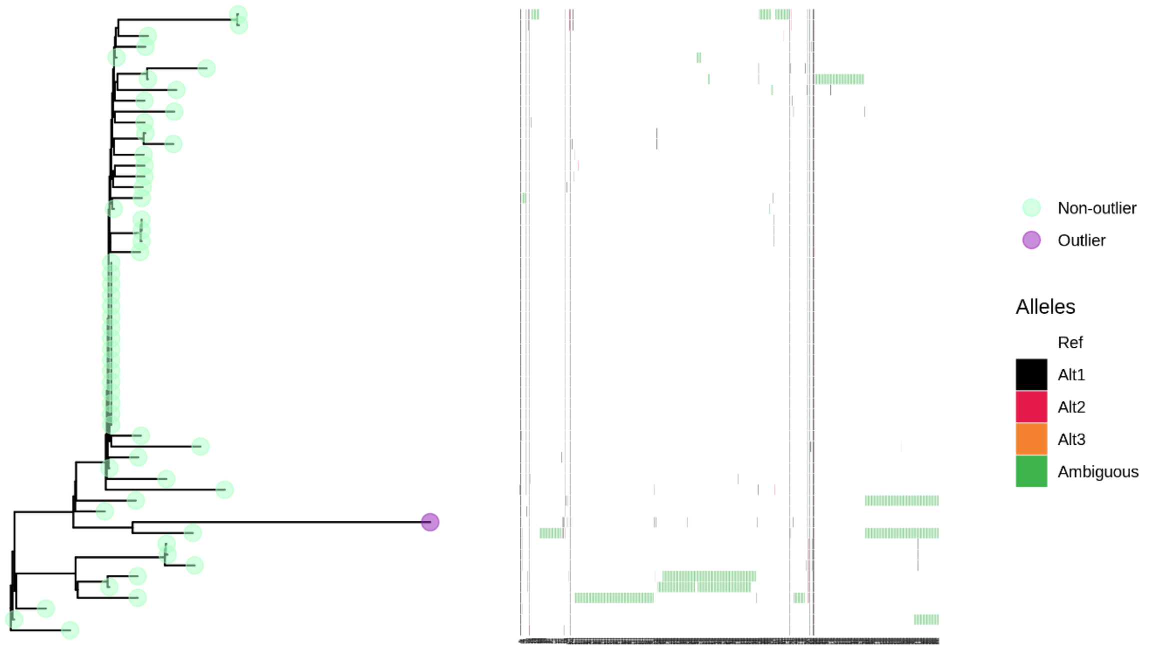

Figure S13

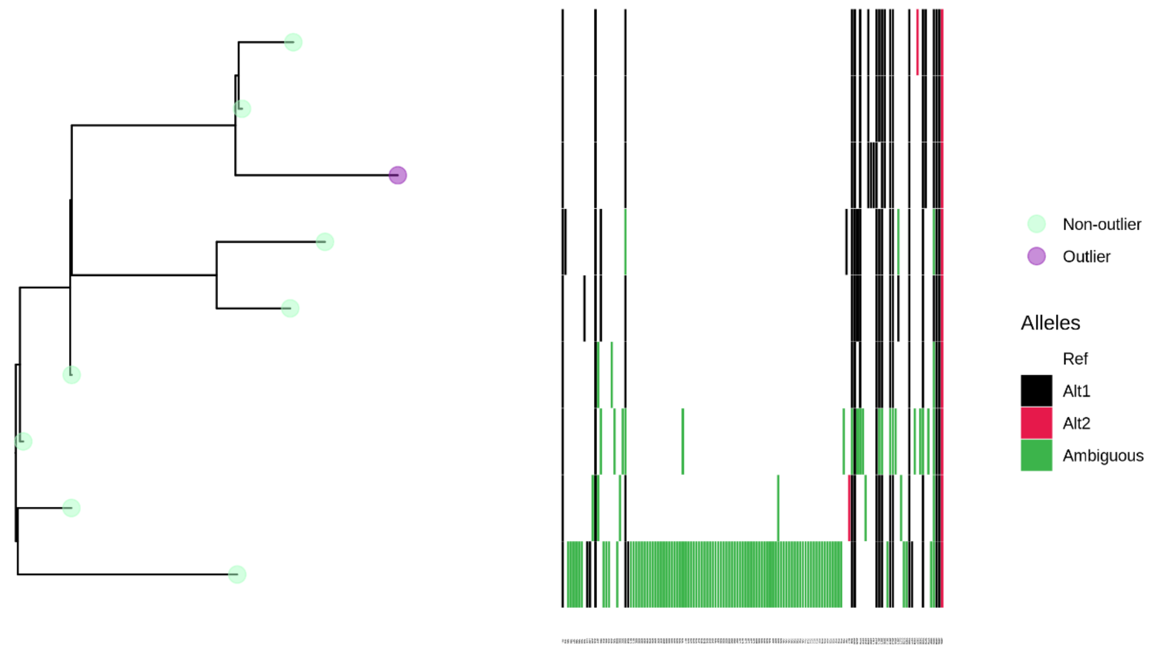


Figure S1-13. Left panels: subclades of the maximum likelihood global SARS-CoV-2 tree flagging those isolates identified as phylogenetic outliers (purple). Right panels: heatmaps providing all mutations (excluding masked sites, see Methods) relative to Wuhan-Hu-1 identified in the strains represented in the left panel. Columns of the heatmap provide SNP positions, in ascending order. Colours denote the status of the non-reference position(s).

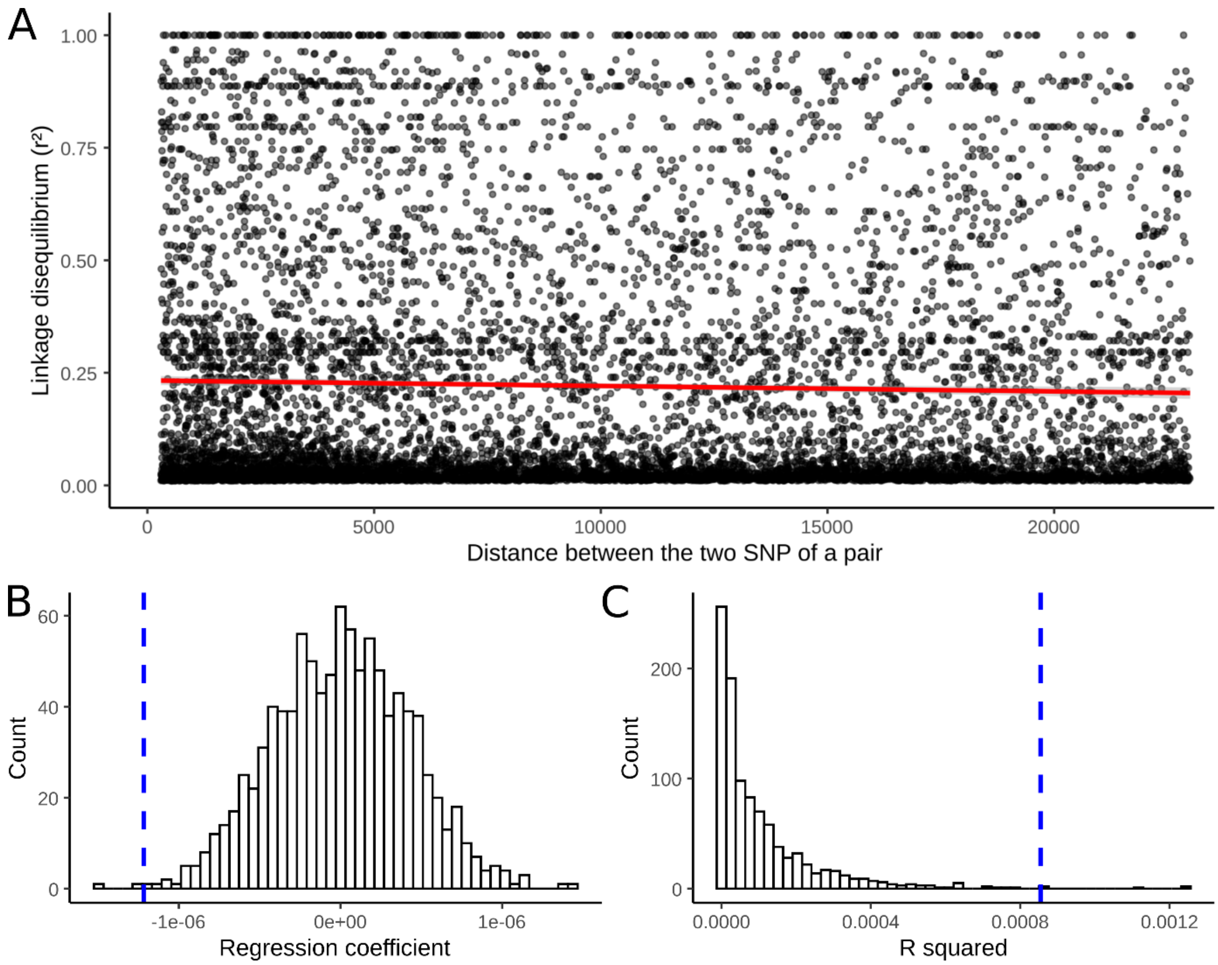


Figure S14: Linkage disequilibrium as a function of distance in the MERS-CoV SNP alignment after discarding C to T mutations. Pairs comprising SNPs differing by a frequency >=0.1 have been discarded, lowering the number of pairs from 9026 to 6066. (A) Linkage disequilibrium (y-axis) for all pairs of SNPs is represented as a function of the distance separating the SNPs of each pair. Red line: fitted linear model (regression coefficient: 1.22e-6; R-squared: 8.56e-4). (B) Distribution of the regression coefficients of the linear models obtained following consideration of 1000 position permuted datasets. Blue dashed line: value of the MERS-CoV true alignment. (C) Distribution of the R squared values of the linear models of 1000 position permuted datasets. Blue dashed line: value of the MERS-CoV true alignment.

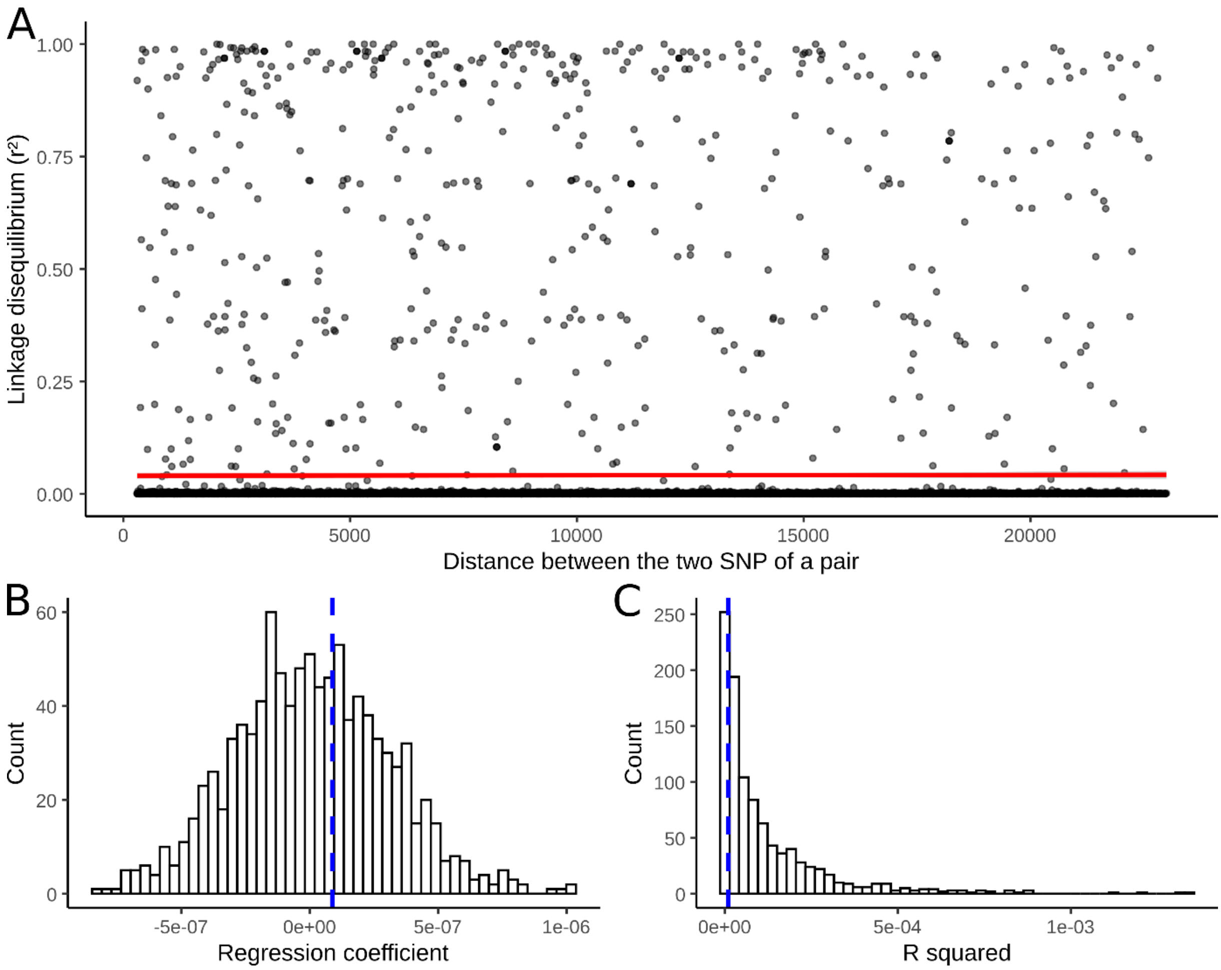


Figure S15: Linkage disequilibrium as a function of distance in the SARS-CoV-2 SNP alignment. Pairs comprising SNPs differing by a frequency >=0.1 have been discarded, lowering the number of pairs from 9470 to 8156. (A) Linkage disequilibrium (y-axis) for all pairs of SNPs is represented as a function of the distance separating the SNPs of each pair. Red line: fitted linear model (regression coefficient: 8.74e-8; R-squared: 9.89e-6). (B) Distribution of the regression coefficients of the linear models obtained following consideration of 1000 position permuted datasets. Blue dashed line: value of the SARS-CoV-2 true alignment. (C) Distribution of the R squared values of the linear models of 1000 position permuted datasets. Blue dashed line: value of the SARS-CoV-2 true alignment.

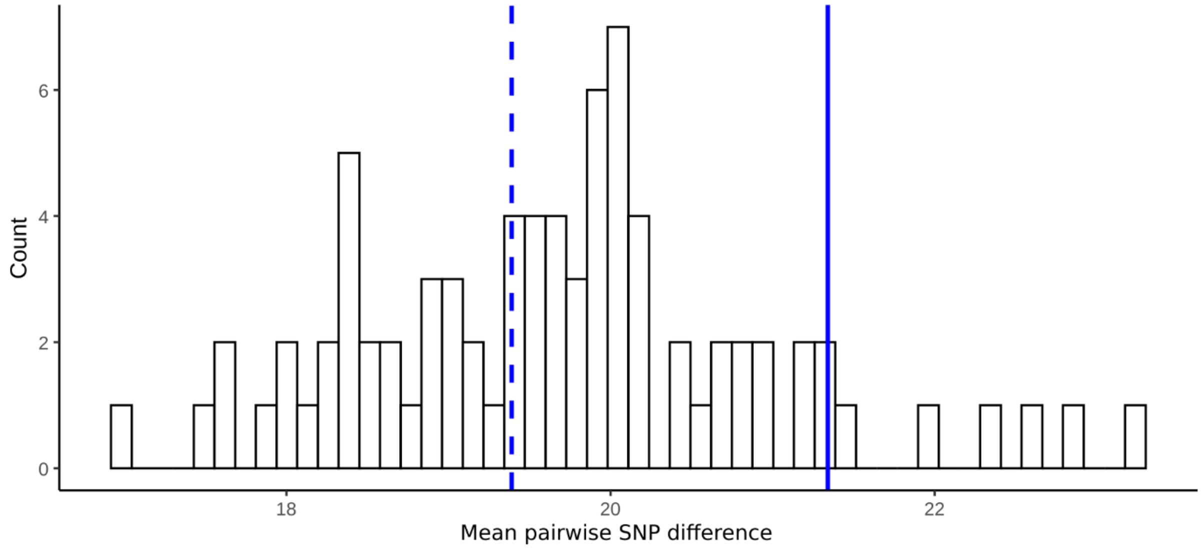


Figure S16: Mean pairwise SNP difference values of simulated and real SARS-CoV-2 alignments. Mean pairwise values of all simulated SARS-CoV-2 alignments (10 replicates for each tested recombination rate: 0, 3e-7, 3e-6, 3e-5, 3e-4, 3e-3 and 3e-2 recombination events per genome per viral replication) are represented as a histogram. Mean pairwise values of the unfiltered (blue line) and filtered (dashed blue line) true SARS-CoV-2 alignment comprising 6,546 SARS-CoV-2 isolates sampled during the month of September 2020 are also represented.
